# Polysialylation is a general feature of immune activation

**DOI:** 10.1101/2025.01.10.632476

**Authors:** Sogand Makhsous, Carmanah Hunter, Charmaine van Eeden, Mohammed Osman, Lisa Willis

**Author notes:** Lisa Willis **Email:**.

## Abstract

Sialic acids are critical regulators of immune responses and the sialic acid-Siglec axis has receive much attention recently for it role as a new immune checkpoint. While α2,3- and α2,6-linked sialosides have been well studied, much less is known about the α2,8-linked polysialic acid (polySia), especially in the adaptive immune system. We demonstrate here that polySia is actually found in all major classes of circulating immune cells, including B and T cells, where it is strongly associated with prior antigen exposure. Cell surface expression of polySia could be observed within 24 h of activation of naïve T cells but did not correlate with expression of the immune-specific polysialyltransferase ST8Sia4. Finally, we assessed the effects of smoking on polySia levels, and noted substantial increased levels of polySia, particularly on effector T cells and preferentially in males.

**Significance Statement:** The immune system is central to human health and holds the key to treating many chronic diseases. We show for the first time that the immunomodulatory glycan polysialic acid is widespread in the human immune system and associated with immune cell activation. Moreover, sex-dependent increases in polysialic acid could be observed on key populations of immune effector cells in smokers. This study offers insight into how glycans might contribute to sex differences in development and progression of chronic diseases.

## Introduction

The immune system is a network of cells that constantly surveil their environment, poised to respond to myriad threats. Resolution of the immune response once the threat is eliminated is critical for maintaining homeostasis. Sustained disruption to this balance leads to the development of chronic diseases, including cancers and inflammatory disorders. Often, sustained immune activation is promoted by environmental factors (e.g., chronic infections, smoking) which eventually result in immune dysregulatory states. Among the known discreet triggers that are known to promote immune dysfunction, smoking is by far one of the most established ones and as such is a primary risk factor for numerous chronic inflammatory diseases (1).

Immune homeostasis is maintained via a network of “checks-and-balances” provided by numerous intracellular and extracellular molecules. This is important as the adaptability of immune cells to their environment is a driver of both normal functions related to infections (e.g. elimination of pathogens), but also may promote the development of maladaptive mechanisms that lead to autoimmunity (2-4). Glycans are critical players that regulate immune responses especially cell surface glycans, which are ideally positioned to modulate the interactions between immune cells and their environments (5-8). Sialic acids in particular represent the front lines of these interactions, given their outer most position at the exposed end of the glycan and capacity to modulate critical receptor-ligand interactions (9,10). Sialic acid-binding immunoglobulin-like lectins (siglecs) are sialic acid receptors that are primarily found on immune cells and control numerous facets of immune activation (11). The sialic acid-siglec axis has recently received widespread attention for its function as immune checkpoints that represent viable therapeutic targets for next generation treatments in cancer and neurodegenerative disorders (12-14). Thus, a better understanding of sialic acids on immune cells will provide added insights related to their fundamental roles in regulating immune cell functions.

One type of sialic acid that is often overlooked is polysialic acid (polySia). PolySia is a long homopolymer of α2,8-linked sialic acids that differs from the widely expressed α2,3- and α2,6-linked monosialosides in that it is mostly limited to the nervous, immune, and reproductive systems of healthy human adults and has only ever been observed to modify a handful of proteins (15). PolySia has long been known to promote cell migration. It is required for neurite outgrowth, axonal migration, and dendritic cell chemotaxis. More recently, it has become apparent that polySia also attenuates immune responses, and thus may represent another dimension of the sialic acid-siglec immune checkpoint axis (16). PolySia inhibited phagocytosis in activated macrophages (17-19), attenuated the ability of dendritic cells to activate T cells (20), and was anti-inflammatory in humanized models of disease (21,22). Curiously, there is no mouse homolog of any known human polySia-binding Siglec (e.g. Siglecs-11 and -16) (11) and there are differences in sequence and expression of polySia-specific Siglecs even between primates (23) suggesting a unique role for polySia in the human immune system that bears further investigation. This is highly relevant as human immune responses and diseases are often not fully captured by rodent models (24,25).

In this study, we demonstrate for the first time that polySia is widespread in all human immune cell types in circulation, including B and T cells. PolySia expression in lymphocytes was strongly associated with prior antigen exposure. Cell surface expression of polySia could be observed within 24 h of activation of naïve T cells but did not correlate with expression of the immune-specific polysialyltransferase ST8Sia4. Finally, we assessed the effects of smoking on polySia levels, and noted substantial increased levels of polySia, particularly on effector T cells and preferentially in males.

## Results

### PolySia is expressed among major human immune cell populations in circulation

Previous attempts to characterize polySia on immune cells were limited by less powerful technology with low dynamic ranges, which generated inconsistent data (20,26). With the development of more powerful lasers and spectral flow cytometers, it is now possible to perform an extensive analysis from a single sample (27). We applied this technology to resolve the question of where polySia is present in circulating immune cells. Peripheral blood mononuclear cells (PBMCs) from 16 healthy human donors (8 male, 8 female) were analyzed by spectral flow cytometry using the polySia lectin GFP-EndoN_DM_ (28) to detect polySia (Fig. 1, S1). To control for the specificity of the polySia signal, all samples were treated in parallel with the polySia hydrolase (Fig. 1A). As expected, we observed polySia^+^ populations in all innate immune cell types, consistent with previous studies (Fig. 1B) (20). We also detected abundant polySia on all NK and NKT cells which is not surprising as human NK/NKT cells express the highly polysialylated protein NCAM/CD56 (29). Within cells of the adaptive immune system, we were intrigued to find both B and T cells contained polySia^+^ populations. We had previously observed polySia in immunoblots of bulk CD3^+^ T cells activated *in vitro* (30) but this analysis represents the first report of polySia^+^ B and T cells in human PBMCs. To determine the relative amounts of polySia on the different cell types, we calculated the median fluorescence intensity of the various polySia^+^ leukocyte populations (Fig. 1C). PolySia levels on NK cells were more than an order of magnitude higher than any other cell type, possibly explaining why polysialylation of other cell types was initially undetected. PolySia^+^ T cells has similar fluorescence intensities as dendritic cells. Surprisingly, even though only ∼10% of B cells were polysialylated, the fluorescence intensity of polySia was substantially higher than that present on monocytes. To provide further insight into major cell types, we showed that NK^bright^ have substantially more polySia per cell than NK^dim^, as expected given that bright/dim refer to NCAM/CD56 expression (Fig. 1D,E). We also observed that classical and non-classical monocytes have similar proportions of polySia^+^ cells but that non-classical monocytes have higher levels of polySia/cell (Fig. 1 F,G). Importantly, no sex-based differences in polySia expression were observed across PBMC populations from healthy donors (Fig. S2). Together, these data demonstrate that polySia is broadly expressed across diverse human immune cell populations in PBMCs, with distinct levels of expression across cell types.

**Figure 1.**
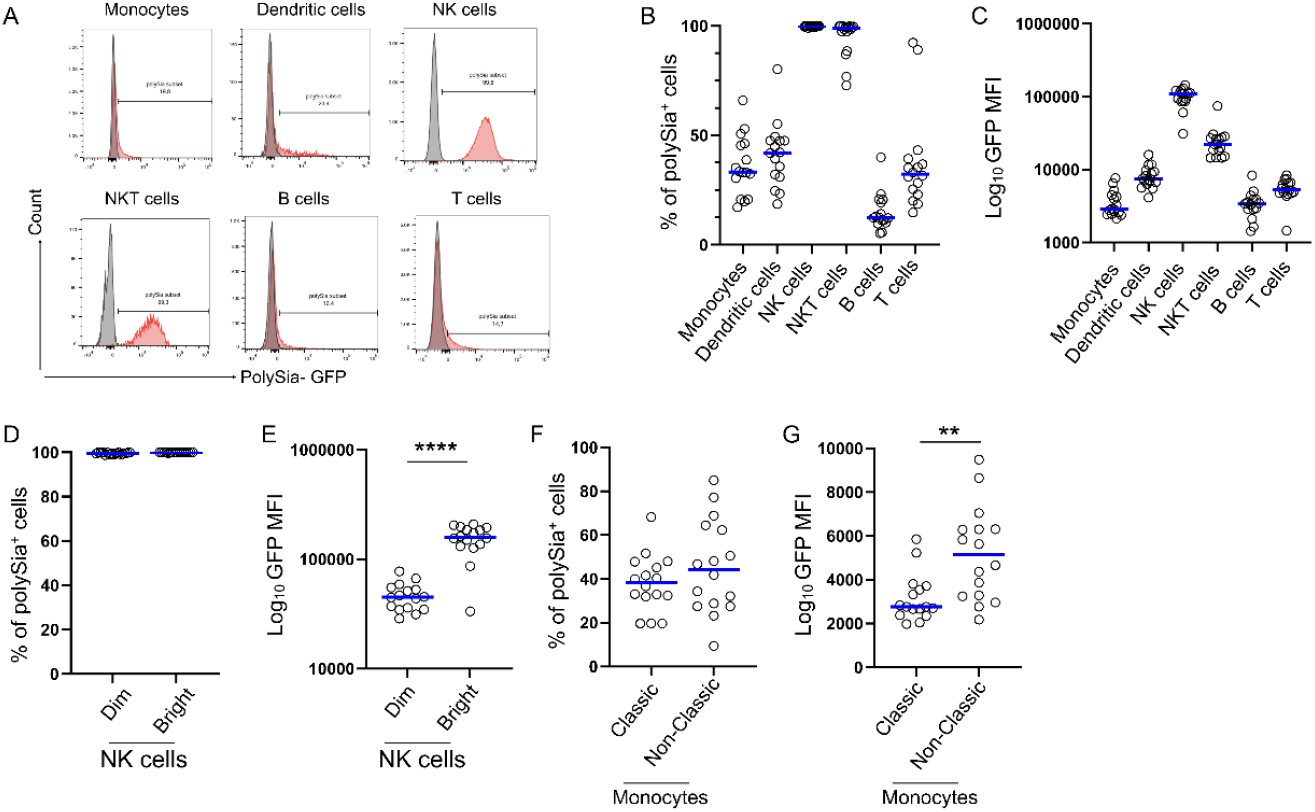
Distribution of polySia among immune cell populations in human PBMCs. A) PBMCs from healthy human adults (20 – 40 yrs old) were analyzed by spectral flow cytometry. GFP-EndoN_DM_ was used to detect polySia. All samples were pre-treated with either a heat-inactivated polySia hydrolase (red) or active polySia hydrolase (grey) in parallel to determine the specificity of the signal. Quantification of the percentage of polySia^+^ cells in each major immune cell population (B) and selected sub-populations (D,F). Relative intensity (log_10_) of median GFP fluorescence intensity (MFI) in polySia^+^ cells representing cell surface polySia expression in major (C) and selected minor (E, G) immune cell populations. Horizontal blue lines represent the median. Statistical analysis of differences between NK and monocyte subpopulations (D-G) was performed using the Mann-Whitney test. (D) ****P < 0.0001; (F) **P = 0.0017.

### PolySia is associated with immune activation in B and T cells

B and T cells are highly heterogeneous polyclonal populations of cells with distinct distributions, functions, and life-spans. Activation of naïve B and T cells promotes their differentiation into memory and effector subtypes. To explore the relationship between immune activation and polySia, we quantified polySia on B and T cells from healthy human PBMCs (Fig. 2, S3,S4). For B cells, naïve cells exhibited the lowest polySia^+^ populations and lowest overall polySia expression, with progressively higher levels as the cells differentiate into antibody-secreting B cells (ASBs) (Fig. 2A-C). For T cells, this trend was even more extreme (Fig. 2D-F). Naïve CD4^+^ and CD8^+^ T cells were almost entirely devoid of surface polySia, while significant populations of antigen experienced cells were highly polysialylated. This polysialylation correlated with the degree of differentiation in memory cells, with T_EM_ having higher proportions of polySia^+^ cells than T_CM_ in both CD4^+^ and CD8^+^ populations. In contrast, the proportion of polysialylated CD8^+^ T_EFF_ was diminished compared to the CD8^+^ T_EM_ population. We observed significant heterogeneity within some subpopulations of T cells, especially those that are more highly differentiated where two maxima appeared, a polySia^high^ and a polySia^lo/-^ population. This was especially apparent in CD8^+^ cells, where the increased amount of polySia^lo/-^ T_EFF_ cells compared to T_EM_ cells suggests that polySia is being lost from some cells.

**Figure 2.**
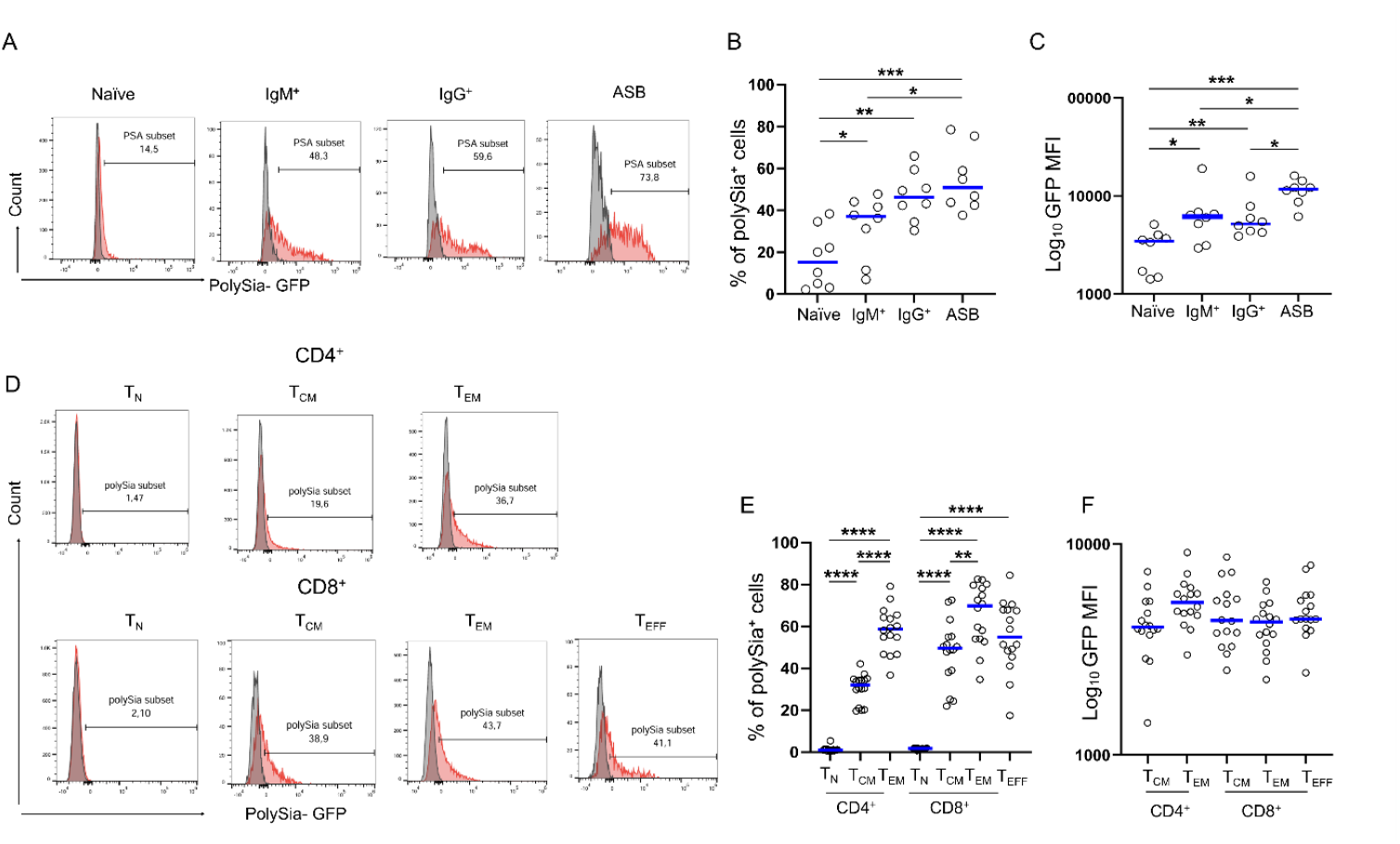
PolySia is associated with activation in B and T cell subtypes. PBMCs from healthy human adults were analyzed by spectral flow cytometry, with a specific focus on B cell (A) and T cell (C) sub-populations. GFP-EndoN_DM_ was used to detect polySia. All samples were pre-treated with either a heat-inactivated polySia hydrolase (red) or active polySia hydrolase (grey) in parallel to determine the specificity of the signal. Quantification of the percentage of polySia^+^ cells (B,E) and the relative intensity of polySia signal in polySia^+^ cells (C,F). PolySia on naïve T cells was virtually undetectable. Comparisons between groups were made using the Mann-Whitney test. Horizontal lines indicate medians. *P < 0.05, **P < 0.01, ***P<0.001, ****P < 0.0001.

Given the profound impact polySia has on cell biology and the substantial increase in polySia^+^ antigen-experienced T cells, we next wanted to determine the temporal dynamics of polysialylation. Naïve CD4^+^ T cells were isolated from healthy PBMCs and activated *in vitro* using α-CD2/3/28 in the presence of IL-2 before analysis using flow cytometry (Fig. 3A-C). Both the proportion of polySia^+^ cells and the intensity of polySia staining increased over several days post-activation. This increase was slow at first, with little change at 24 h, suggesting that polysialylation is not an immediate response to antigen stimulation. We hypothesized that the delay in polysialylation might be due to the requirement for synthesis of the polysialyltransferase responsible for making polySia in immune cells, ST8Sia4 (19,20,31,32). We analyzed the mRNA levels of *ST8SIA4* by qPCR and found no statistically significant differences (Fig. 3D), suggesting that polysialylation is controlled by alternative levels of regulation.

**Figure 3.**
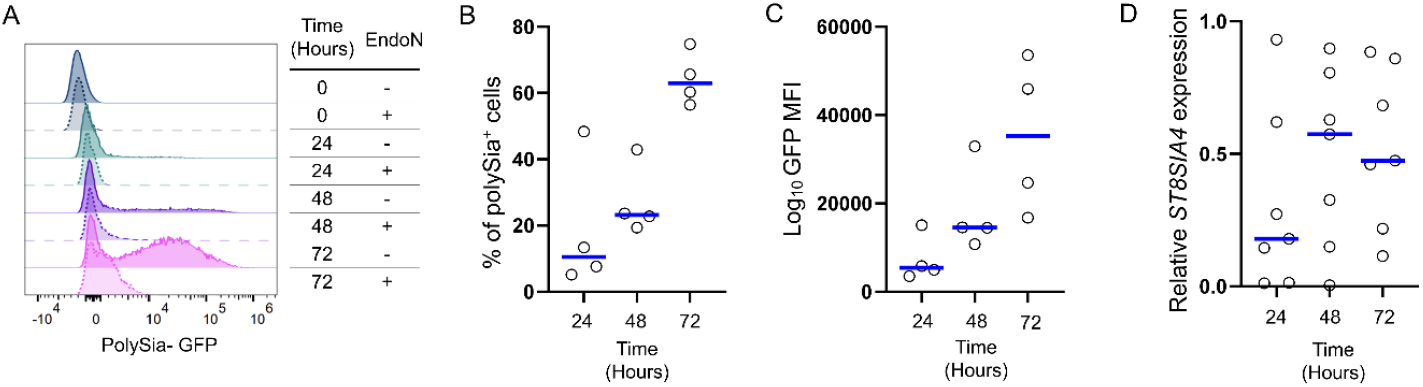
PolySia is expressed after activation of naïve CD4^+^ T cells but is not correlated with ST8Sia4 mRNA levels. Naïve CD4+ T cells were isolated from healthy PBMCs and activated *in vitro*, after which samples were taken at 24, 48, and 72 hr. PolySia expression was analyzed by flow cytometry (A), and changes in the percentage of polySia^+^ cells (B) and GFP median fluorescence intensity (MFI) (C) were measured. D) *ST8SIA4* mRNA expression was measured by qPCR using 18S rRNA as a standard.

### PolySia expression is higher in smokers, especially in males

Smoking is a well-known contributor to chronic inflammation, with previous studies showing that smokers have elevated levels of systemic inflammatory markers such as C-reactive protein (CRP) and fibronectin, and are at higher risk for autoimmune diseases and cancer (33). To explore the impact of smoking-induced chronic inflammation on polySia expression in immune cells, we compared polySia levels across different cell populations in smokers (Fig. 4). PolySia expression was generally higher in smokers, with a significant increase observed in the proportion of polySia^+^ cells in the monocyte population (Fig. 4A). Among monocyte subpopulations, classical monocytes exhibited a significantly higher proportion of polysialylation (Fig. 4B) though no change in the total amount of polySia/cell. While the proportion of polySia^+^ CD3^+^ T cells were not statistically higher in smokers (p = 0.17), the strong trend led us to examine T cell subpopulations in more detail. Intriguingly, no differences were observed in the CD4^+^ compartment but smokers had higher levels of polySia in the CD8^+^ compartment (Fig. 4C). These differences become significant for both measured metrics as the CD8^+^ cells further differentiate to T_EFF_. Moreover, we observed striking sex differences in both monocyte and CD8^+^ T_EFF_ cells (Fig. 4D-F), with males typically having higher proportions of polySia^+^ and also higher levels of polySia/cell. In summary, these data indicate that smokers exhibit higher polySia expression across various immune cell populations, particularly in monocytes and CD8^+^ T cells, with polySia potentially contributing to sex differences in chronic inflammation.

**Figure 4.**
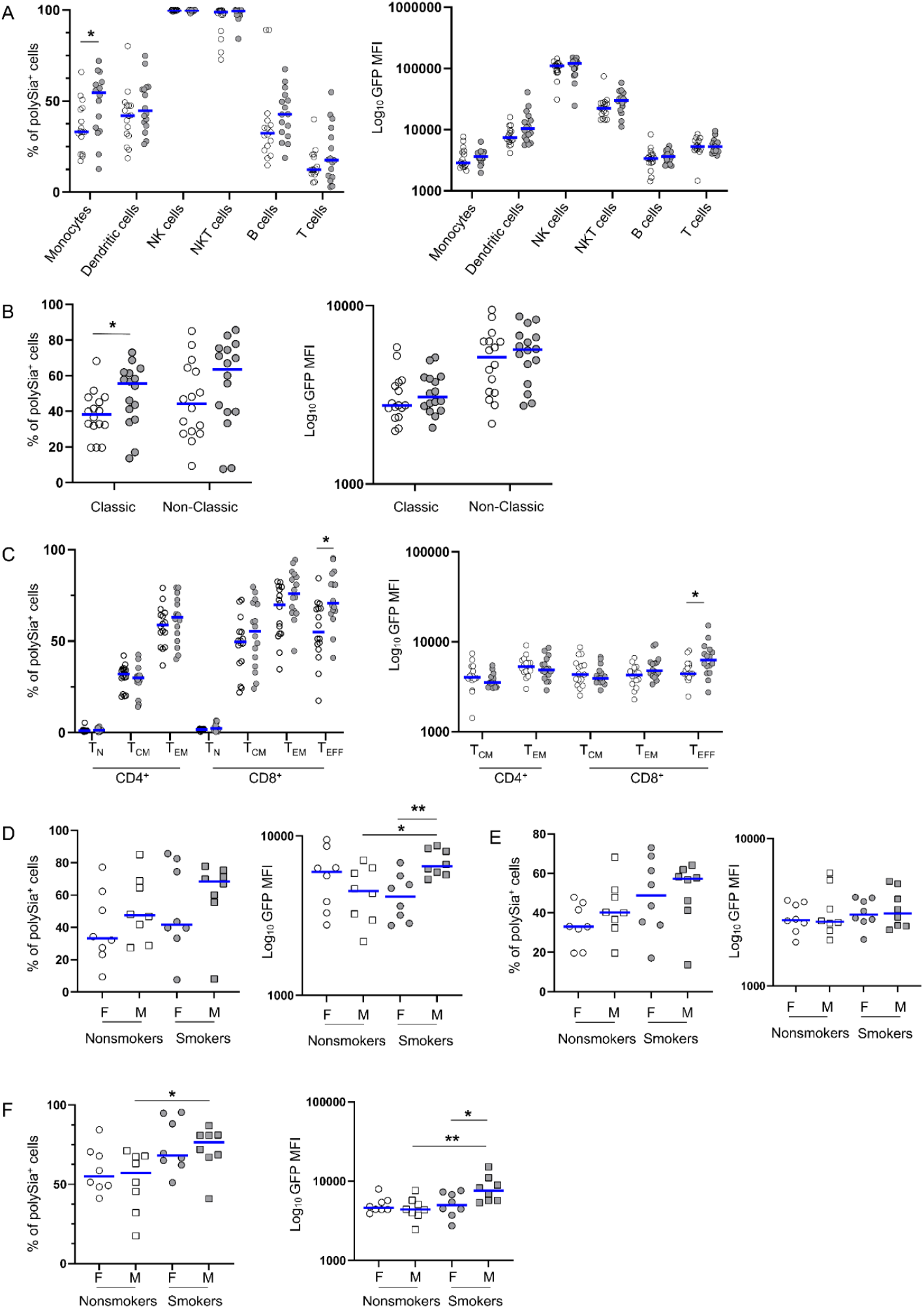
PolySia levels are increased in smokers with some sex-specific differences. PBMCs from otherwise-healthy human smokers and non-smokers (20 – 40 yrs old) were analyzed by spectral flow cytometry. GFP-EndoN_DM_ was used to detect polySia. All samples were pre-treated with either a heat-inactivated polySia hydrolase or active polySia hydrolase in parallel to ensure the specificity of the signal. Quantification of the percentage of polySia^+^ cells and the relative median intensity of staining in polySia^+^ cell were measured for major immune populations (A), monocyte subpopulations (B), and T cell subpopulations (C). This data is shown disaggregated by sex for samples with significant differences – total monocytes (D), classic monocytes (E), and CD8^+^ T_EFF_ cells (F). For A-C, non-smokers are in open circles (◯), smokers in grey circles 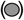. Comparisons between groups were made using the Mann-Whitney test. Horizontal lines indicate medians. *P < 0.05, **P < 0.01.

## Discussion

The role of polySia in the human immune system is critically underexplored. Here, we performed an in-depth analysis of cell surface polySia on PBMCs from healthy young adults and smokers and show that it is widespread across all major immune cell populations, including those previously thought to lack polySia. PolySia is also clearly associated with immune activation in multiple different cell types. This phenomenon was first described in activated dendritic cells (20,34) and now also holds true for B and T cells. The significant differences in polySia levels across different T cell subsets is particularly striking and begs the question of how polySia may influence T cell biology.

PolySia is most strongly associated with a migratory phenotype and thus may influence T cell migration. PolySia is required for neurite outgrowth and axonal migration during brain development (15,35), as well as cytotrophoblast migration during formation of the placenta (36). The glycan has also been shown to be required for migration of CCR7^+^ dendritic cells towards its ligand CCL21 (32,34,37,38). At roughly 70%, T cells are the most abundant white blood cell in circulation and their migration patterns are highly dependent on their activation state. T_N_ and T_CM_ cells circulate constantly between the blood, lymphatics, and secondary lymphoid organs, whereas T_EM_ cells surveil blood and non-lymphoid tissue (39). Ironically, the cell types with the lowest proportion of polySia^+^ cells – T_N_ and T_CM_ – migrate to secondary lymphoid organs through interactions between CCR7 and CCL21, the chemokine-ligand pair that are highly dependent on polySia for migration of dendritic cells. If polySia does indeed stimulate the migration of T cells, one possible explanation could be that polySia on T_CM_ cells increases their association with secondary lymphoid organs so that they circulate more frequently in search of a cognate antigen. Similarly for the highly polysialylated T_EM_ cells, the presence of polySia may enhance their migration towards non-lymphoid tissues, depending on the particular chemokine/receptor that is modified.

PolySia, like other sialic acids, is emerging as an immune checkpoint (16,40,41) and thus may also influence T cell activation. In certain contexts, polySia can attenuate immune responses, inhibit the release of proinflammatory cytokines, and decreases phagocytosis (17,19,20), though the mechanisms supporting these phenotypes remain mostly unknown. The dynamic nature of polySia, where it increases over the course of several days upon activation of naïve T cells and then is lower in T_EFF_ cells, could hint at a glycan that attenuates T cell activity, although no sialic acid receptor has ever been observed on T cells (42). Clearly more research is required to understand the role of polySia in T cell biology. The lack of association between mRNA expression of the polysialyltransferase ST8Sia4 and cell surface polySia levels in T cells is not uncommon for sialyltransferases, which are sometimes subject to additional levels of regulation (43). Glycosylation itself is one such mechanism of alternative regulation that is used extensively in T cells, as in the case of PD-1 (44). Understanding these regulatory pathways will be crucial for elucidating the broader functional roles of polySia in immune modulation.

We were intrigued to discover that polysialylation is higher in smokers, particularly in the monocyte and CD8^+^ T cell populations. Smoking induces chronic inflammation and smokers are much more likely to develop numerous chronic diseases. We and others have shown that polySia is increased in cancer and autoimmune diseases (15,45). All of the donor PBMCs analyzed in this study came from young adults (20 – 40 years of age), suggesting that changes in polySia occur relatively early and may have profound effects later in life. Other cell populations, like B cells, trended towards higher levels of polySia and it would be interesting to know if that increase became significant with longer or heavier smoking. The lack of sex differences in most cell types surprised us as we previously showed that males have higher levels of polySia in serum (46). However, we had not controlled for smoking in that study and gender norms might suggest that more of the samples we obtained from males were smokers. The sex differences in polySia on CD8^+^ T_EFF_ cells has potential implications for development of chronic diseases, especially if polySia attenuates the activity of these cytotoxic lymphocytes, as they are responsible for eliminating pathogens and cancerous cells and are well established to be less effective in males compared to females (47).

To conclude, our work identifies polySia as a dynamically regulated glycan with broad implications for human immune cell function. The selective expression of polySia in activated immune cells and its upregulation in chronic inflammation underscore its potential as both a biomarker and a therapeutic target. Further studies are warranted to elucidate the precise mechanisms by which polySia modulates immune activity and to explore the role of polySia in the context of inflammatory diseases such as cancer and autoimmunity. This work will provide a critical platform to investigate the role of polySia in regulating the immune response in health and disease.

## Materials and Methods

### PBMCs

Cryopreserved PBMCs were obtained from STEMCELL Technologies (Table S1). Cells were thawed in a 37°C water bath for approximately 2 min, followed by washing with Dulbecco’s phosphate-buffered saline (DPBS). After washing, cells were centrifuged at 180 × *g* for 10 min at room temperature to prepare for downstream applications. Cell count and viability were assessed using a hemocytometer and trypan blue exclusion.

### CD4^+^ naïve T cell isolation and manipulation

Naïve CD4^+^ T cells were purified by negative selection using the EasySep™ Human Naïve CD4^+^ T Cell Isolation Kit II (STEMCELL Technologies) following the manufacturer’s instructions. Briefly, washed PBMCs were resuspended at a concentration of 5 × 10^7^ cells/mL in DPBS supplemented with 1 mM EDTA. Cells were incubated with the isolation cocktail (50 µL/mL) for 5 minutes, followed by the addition of RapidSpheres™ (50 µL/mL). The sample volume was adjusted to 2.5 mL with DPBS containing 1 mM EDTA. The tube was then placed in the EasySep™ Magnet for 3 min, and unlabeled cells were collected in a fresh tube.

Isolated naïve CD4^+^ T cells were cultured in ImmunoCult™-XF T Cell Expansion Medium (STEMCELL Technologies) supplemented with 25 U/mL IL-2 (Miltenyi Biotec) at a concentration of 1 × 10^6^ cells/mL. Cells were activated using ImmunoCult™ Human CD3/CD28/CD2 T Cell Activator (STEMCELL Technologies) and incubated at 37 °C with 5% CO_2_ for 3 days. Samples were collected immediately post-isolation and at 24, 48, and 72 hours post-activation. PolySia expression on the cell surface was analyzed by flow cytometry, and ST8Sia4 mRNA levels were quantified via qPCR.

### Flow cytometry

For flow cytometry analysis, cells were incubated with Zombie NIR viability dye (BioLegend) for 10 min at 4°C to label dead cells. Fc receptors were blocked using Human TruStain FcX (BioLegend). To confirm the specificity of the polySia signal, cells were treated with either active or heat-inactivated polySia hydrolase (EndoN) at 10 µg/mL for 30 minutes. EndoN was expressed and purified as previously described (48) and heat-inactivated at 100°C for 10 min. Cells were then stained with antibodies targeting cell-surface markers to identify polySia expression on different populations (Table S2). PolySia was detected using polySia lectin (EndoN_DM_) conjugated to GFP (46), added to the antibody master mix at 10 µg/mL. Staining was performed for 30 minutes on ice, after which cells were washed and resuspended in 300 µL of wash buffer. Samples were analyzed on a Cytek Aurora flow cytometer, and data were processed using FlowJo software (version 10.8.1).

The proportion of polySia^+^ cells in various populations was determined by gating polySia-expressing subpopulations based on EndoN-treated corresponding samples. Cell counts for each population were normalized to the total live cell count in the respective sample. The proportion of polySia^+^ cells in each population was then calculated by dividing the normalized count of polySia^+^ cells by the normalized count of the total population. To normalize the median fluorescent intensity (MFI) of polySia^+^ cells, the GFP MFI value of the polySia^+^ subpopulation in the heat-inactivated EndoN-treated sample was subtracted from the GFP MFI value of the corresponding population in the active EndoN-treated sample.

### *ST8SIA4* qPCR

RNA was extracted from activated T cells (described above), using the RNeasy Mini kit (Qiagen). Subsequently, real-time PCR was carried out using Taqman Fast Virus 1 step Master Mix (Applied Biosystems), and ST8Sia4 (Hs00379924_m1) and 18s rRNA (Hs03928985_g1) gene expression kits (ThermoFisher Scientific), according to manufacturer instructions. ST8Sia4 gene expression was normalized to 18S rRNA. PCRs were conducted in triplicate, and expression calculated as ddCT.

### Statistics

Data analysis was conducted using GraphPad Prism 10. Group comparisons were performed using the Mann-Whitney U test. Statistical significance was denoted as follows: *P* < 0.05 (\**), P < 0*.*01 (**), P < 0*.*001 (\*\*\**), and *P* < 0.0001 (****).

## Acknowledgments

Funding for this work was provided by the Natural Sciences and Engineering Research Council of Canada (Grant/Award Number: RGPIN-2021-02888) and Canadian Glycomics Network (Grant/Award Number: CD-87).

## Supporting information

**Table S1.** Demographic information for donors.

|  |  | Number<br>of samples | Average<br>age | Average BMI<br>(kg/m<sup>2</sup>) |
| --- | --- | --- | --- | --- |
| **Nonsmokers** | Female | 8 | 29.125 | 27.80724 |
|  | Male | 8 | 31.125 | 27.46477 |
| **Smokers** | Female | 8 | 31.5 | 28.84452 |
|  | Male | 8 | 31 | 30.81803 |

**Table S2.** Abtibodies used in this study.

| Panel 1: Total PBMC and T cells |  |  |  |
| --- | --- | --- | --- |
| Specificity | Fluorochrome | Source | Identifier |
| Live/Dead | Zombie NIR | Biolegend | 423105 |
| CD3 | BV510 | Biolegend | 317332 |
| CD4 | BV605 | Biolegend | 344646 |
| CD8 | BV711 | Biolegend | 344734 |
| CD14 | AF700 | Biolegend | 367113 |
| CD16 | BV785 | Biolegend | 302045 |
| CD20 | PE-Dazzle 594 | Biolegend | 302347 |
| CD45RA | PE | Biolegend | 304108 |
| CD56 | BV421 | Biolegend | 302347 |
| CD95 | AF647 | Biolegend | 305618 |
| CD197 | PE-Cy5 | Biolegend | 353272 |
| HLA-DR | PE-Cy7 | Biolegend | 361611 |
| PolySia | GFP | NA | NA |
| Panel 2: B cells |  |  |  |
| Specificity | Fluorochrome | Source | Identifier |
| Live/Dead | Zombie NIR | Biolegend | 423105 |
| CD3 | BV510 | Biolegend | 317332 |
| CD14 | AF700 | Biolegend | 367113 |
| CD19 | BV605 | Biolegend | 302243 |
| CD20 | PE-Dazzle 594 | Biolegend | 302347 |
| CD27 | PE-Cy7 | Biolegend | 356411 |
| CD56 | BV421 | Biolegend | 302347 |
| IgD | PerCP | Biolegend | 348233 |
| IgM | AF647 | Biolegend | 314535 |
| IgG | BV711 | Biolegend | 410739 |
| PolySia | GFP | NA | NA |
| Panel 3: T cells |  |  |  |
| Specificity | Fluorochrome | Source | Identifier |
| Live/Dead | Zombie NIR | Biolegend | 423105 |
| CD3 | BV510 | Biolegend | 317332 |
| CD4 | BV605 | Biolegend | 344646 |
| CD8 | BV711 | Biolegend | 344734 |
| CD45RA | PE | Biolegend | 304108 |
| CD95 | AF647 | Biolegend | 305618 |
| CD197 | PE-Cy5 | Biolegend | 353272 |
| PolySia | GFP | NA | NA |

**Figure S1.**
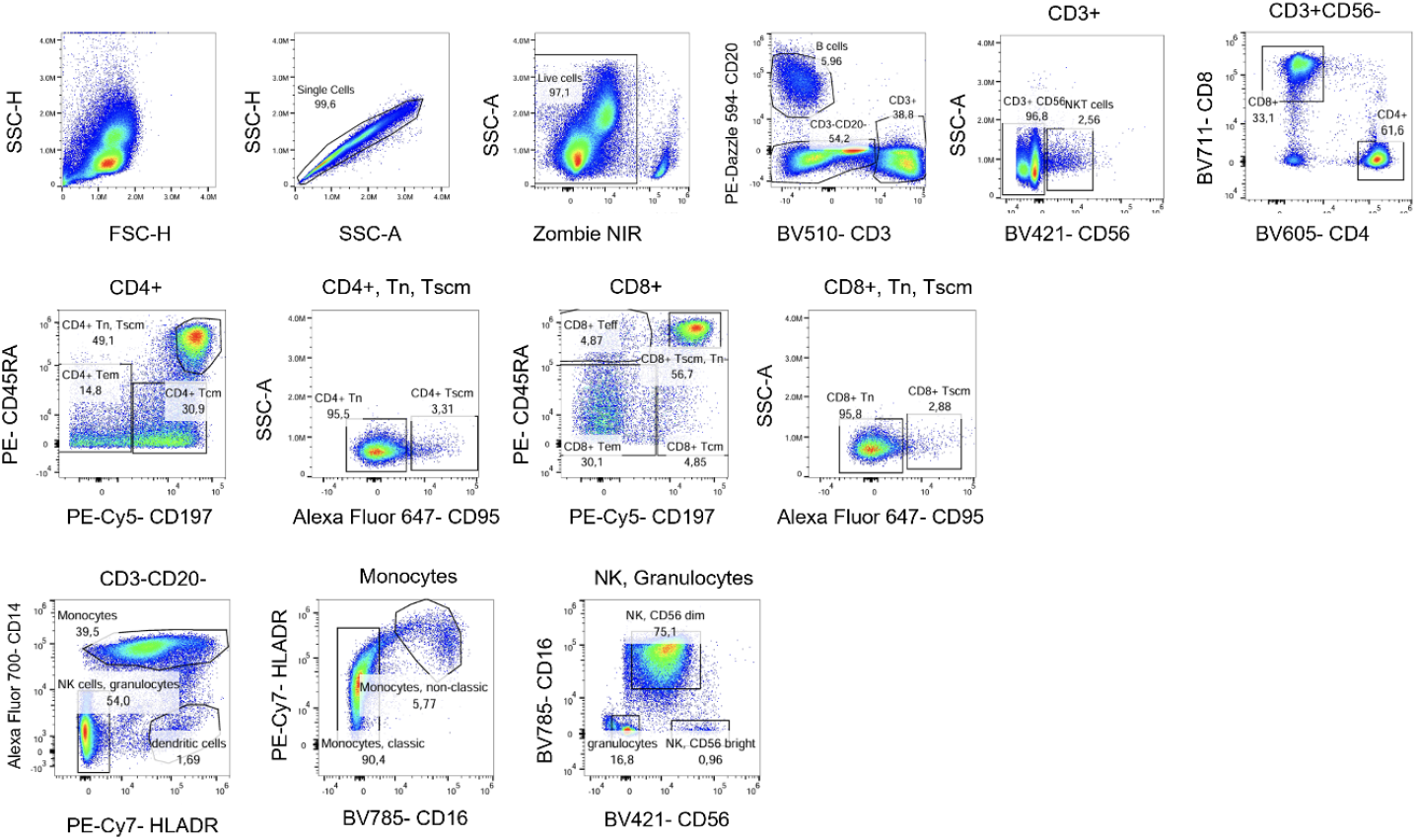
Gating strategy for major and minor immune cell populations from PBMCs (excluding B cell subpopulations).

**Figure S2.**
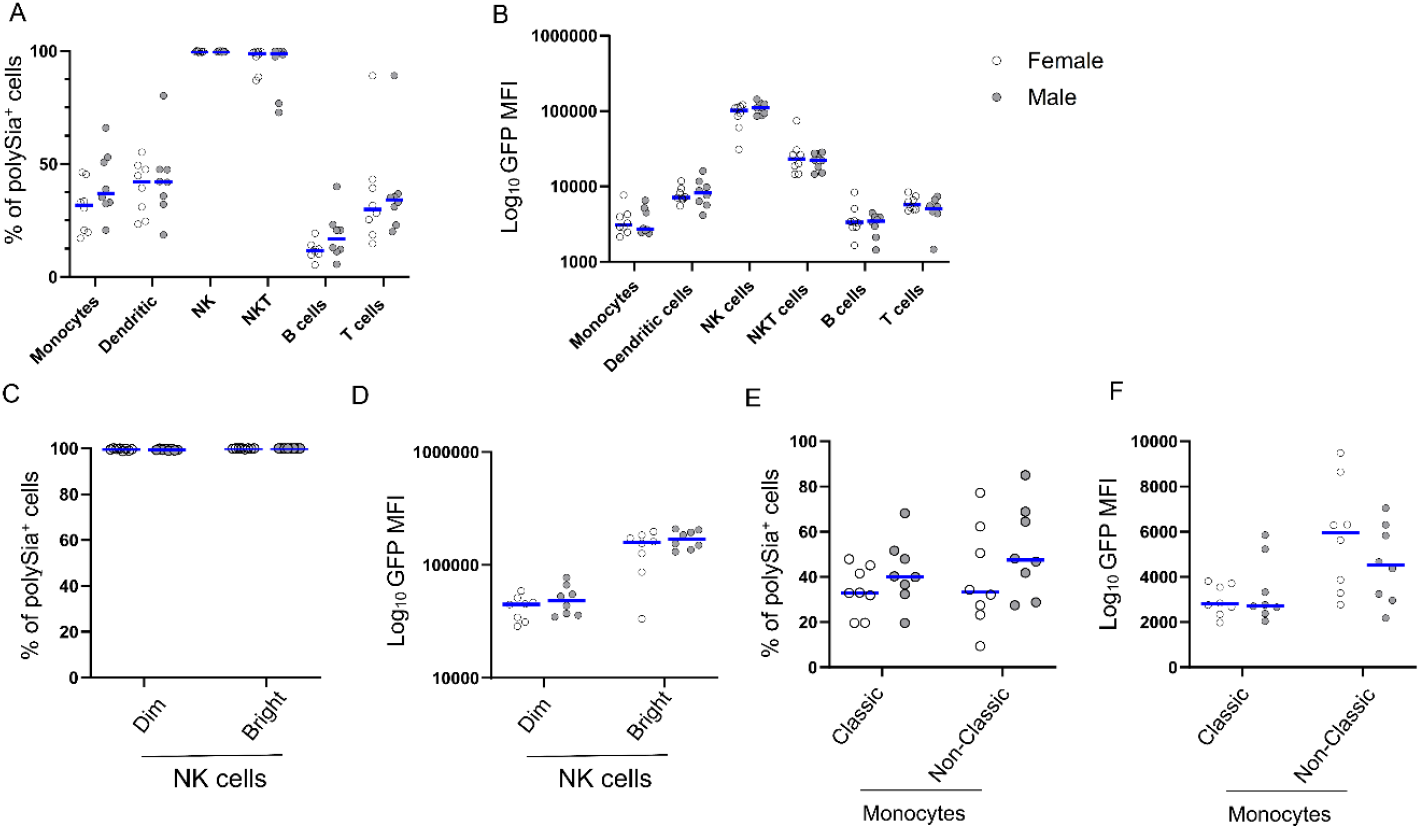
Distribution of polySia among immune cell populations in male and female human PBMCs. Sex disaggregated data for polySia expression among immune cell populations in human PBMCs: The percentage of polySia-expressing cells and the median fluorescence intensity (MFI) of the GFP signal were analyzed for potential sex differences using the Mann-Whitney test. No significant differences in polySia expression were observed between male and female donors.

**Figure S3.**
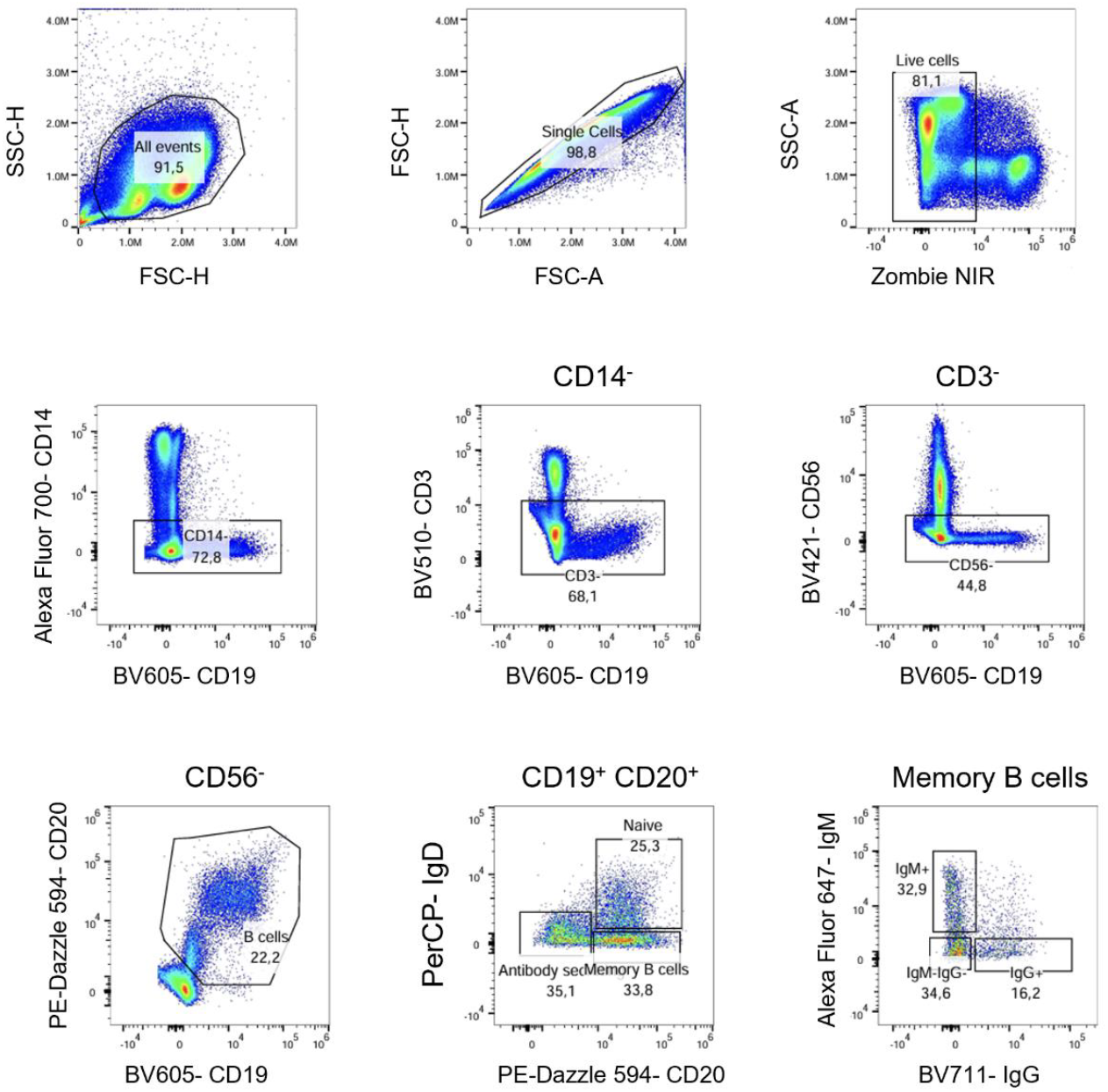
Gating strategy for B cell sub-populations from PBMCs.

**Figure S4.**
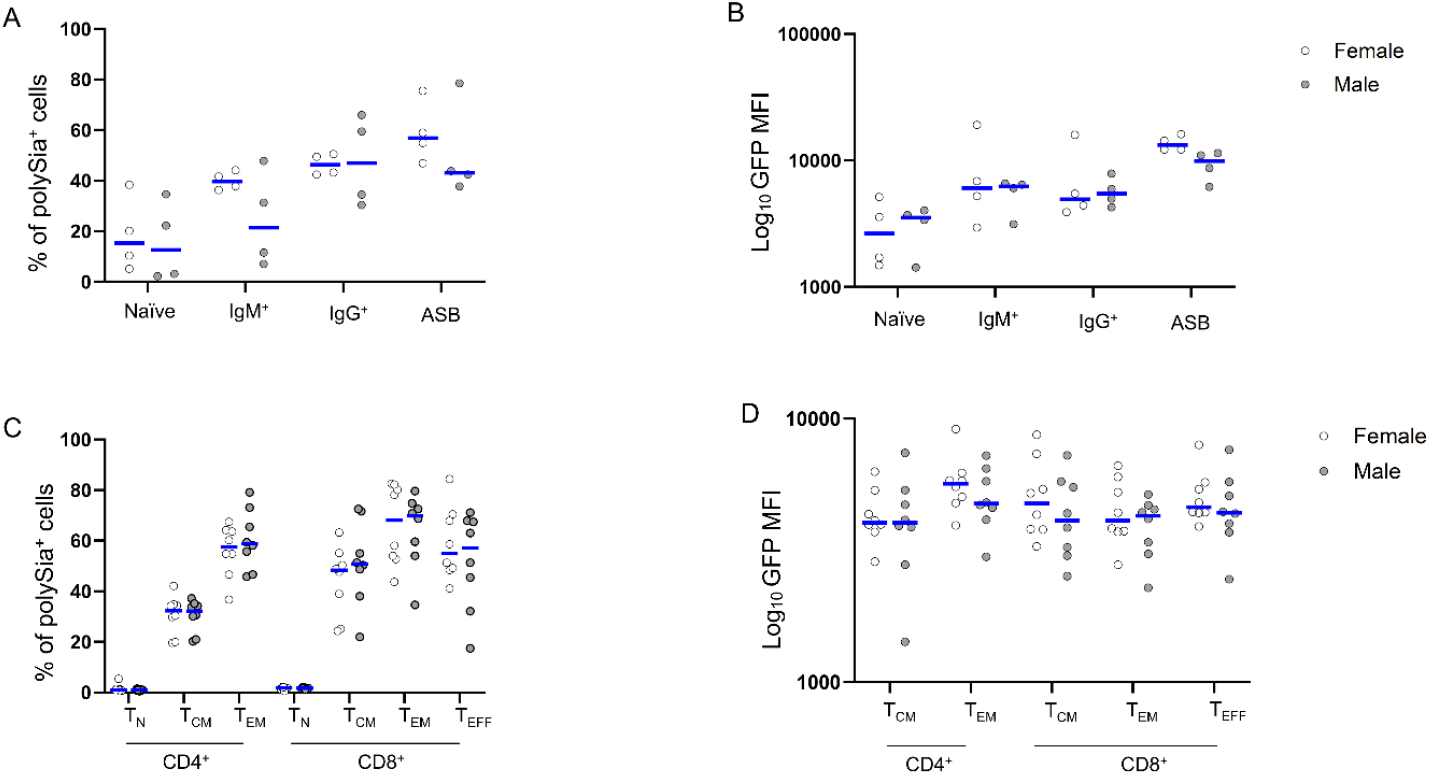
Distribution of polySia among immune cell populations in male and female healthy human PBMCs. Sex disaggregated data for polySia expression in different subpopulations of the adaptive immune system: The percentage of polySia-expressing cells and the median fluorescence intensity (MFI) of the GFP signal were analyzed for potential sex differences using the Mann-Whitney test. No significant differences in polySia expression were observed between male and female donors.

**Figure S5.**
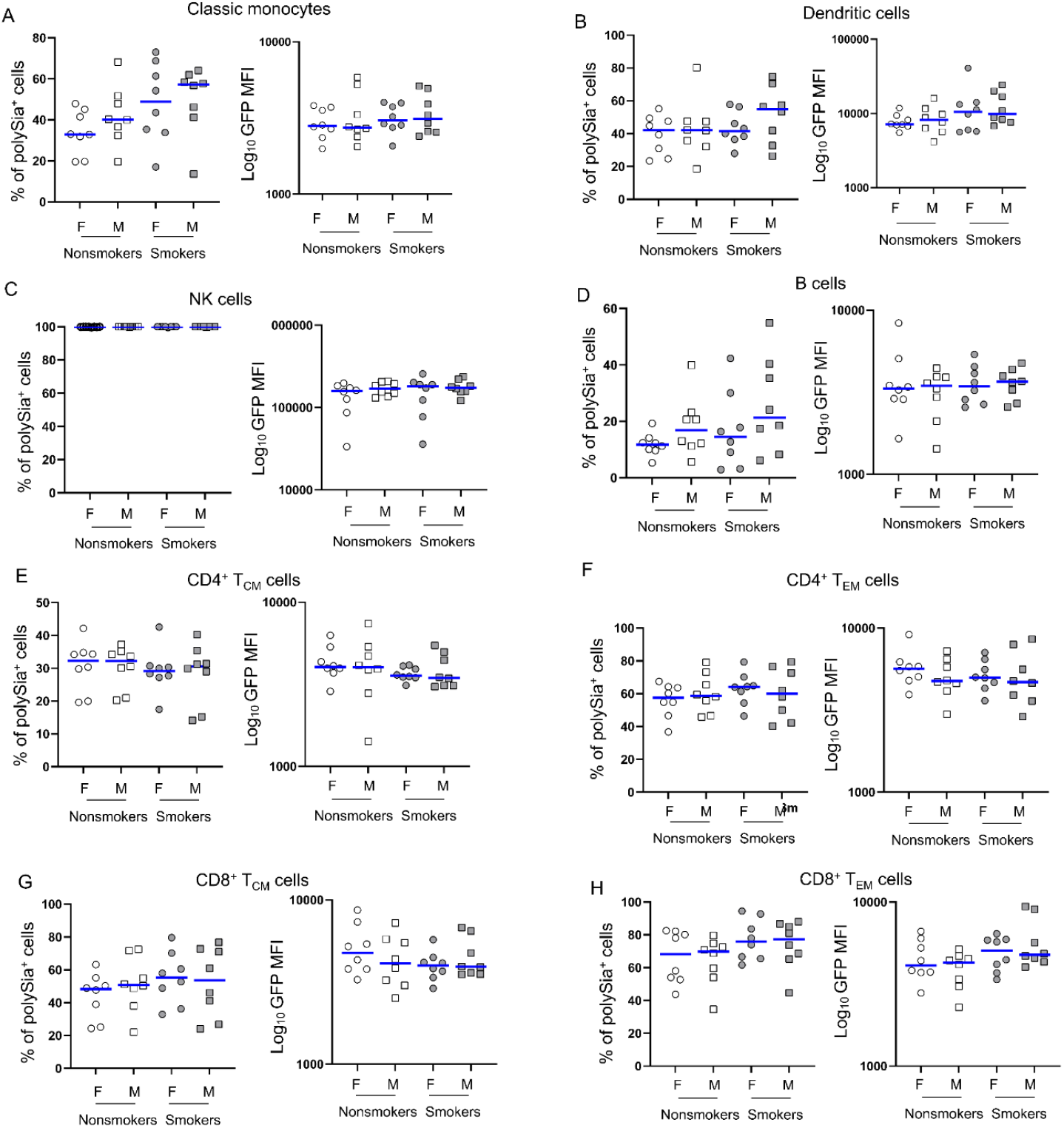
Distribution of polySia among immune cell populations in male and female human PBMCs from smokers and non-smokers. Sex disaggregated data for polySia expression in different subpopulations of the adaptive immune system for populations where no significant differences were observed. The percentage of polySia-expressing cells and the median fluorescence intensity (MFI) of the GFP signal were analyzed for potential sex differences using the Mann-Whitney test.

## Notes

### Competing Interest Statement

The authors have declared no competing interest.

